# Motif V acts as a Regulator of Energy Transduction Between the Flavivirus NS3 ATPase and RNA Binding Cleft

**DOI:** 10.1101/842088

**Authors:** Kelly E. Du Pont, Russell Davidson, Martin McCullagh, Brian J. Geiss

## Abstract

The unwinding of double-stranded RNA intermediates is a critical component for the replication of flavivirus RNA genomes. This function is achieved by the C-terminal helicase domain of nonstructural protein 3 (NS3). As a member of the superfamily 2 (SF2) helicases, NS3 is known to require the binding and hydrolysis of ATP/NTP to translocate along and unwind double-stranded nucleic acids. However, the mechanism of energy transduction between the ATP and RNA binding pockets is not well understood. Previous molecular dynamics simulations published by our group have identified Motif V as a potential “communication hub” for this energy transduction pathway. In order to investigate the role of Motif V in this process, a study combining molecular dynamics, biochemistry and virology has been employed. Mutations of Motif V were tested in both replicon and recombinant protein systems to investigate viral genome replication, RNA binding affinity, ATP hydrolysis activity and helicase unwinding activity. Using these analyses, we found that T407A and S411A in Motif V demonstrated increased turnover rates, suggesting that the mutations causes the helicase to unwind dsRNA more quickly than WT. Additionally, simulations of each mutant were used to probe structural changes within NS3 caused by each mutation. These simulations indicate that Motif V controls communication between the ATP binding pocket and the helical gate. These data help define the linkage between ATP hydrolysis and helicase activity within NS3 and provide insight into the biophysical mechanisms for ATPase driven NS3 helicase function.

---

Arthropod-borne flaviviruses, such as yellow fever virus, Japanese encephalitis virus, Zika virus, West Nile virus (WNV) and dengue virus, are a major health concern in the tropical and subtropical regions of the world (1, 2). Infection from these viruses cause disease symptoms ranging from flu-like illness to encephalitis, hemorrhagic fever, coma, and potentially death (3). Over half of the world population is at risk for infection from one or more of these viruses (4, 5). Dengue virus, specifically, infects around 50 million people each year, and of those individuals, 20,000 contract dengue hemorrhagic fever leading to their mortality (6). Additionally, WNV over the past 20 years within the 48 continental United States has around 50,000 clinical infections with a 5% mortality rate (7–9). Currently, there are no approved antiviral drugs for treating flaviviral infections and the vaccines in circulation, like the yellow fever vaccine, are not readily available worldwide for most flaviviruses (10–12). In order to develop new antiviral treatments (drugs and vaccines) for these viruses, a fundamental understanding of how these flaviviruses replicate is required.

Flaviviruses (*Flaviviridae* family) have a positive-sense single-stranded RNA (ssRNA) genome that is approximately 11 kb in length. The viral RNA genome is translated into a single polyprotein that is subsequently cleaved by host and viral proteases into three structural proteins (C, prM and E) and eight nonstructural proteins (NS1, NS2A, NS2B, NS3, NS4A, 2K, NS4B and NS5) (13, 14). The structural proteins contribute to the formation of mature viral particles, while the nonstructural proteins are responsible for replication of the viral genome and protecting the virus from attacks by the host cell innate immune system (13). Once the viral proteins are post-translationally processed, the positive-sense genomic ssRNA is used as a template to create a negative-sense anti-genomic RNA, forming a double-stranded RNA (dsRNA) intermediate complex (15, 16). The nascent negative-sense ssRNA serves as a template strand for producing new positive strand RNAs that can be packaged into viral particles, translated into new proteins, or interfere with the RNA decay pathway (17, 18). Therefore, the unwinding of dsRNA intermediate to produce a free negative ssRNA for positive strand synthesis is a critical component for the replication of flavivirus RNA genomes (15). This function is achieved by the C-terminal helicase domain of NS3.

Helicases are ubiquitous enzymes that are classified into superfamilies based on their primary structure and highly conserved motifs (19–21). NS3 helicase (NS3h) is a member of superfamily 2 (SF2) helicases (22). The structure of NS3h consists of three subdomains; subdomain 1 and subdomain 2 are RecA-like structures that are highly conserved across SF2 helicases (23). The third subdomain is unique to the viral/DEAH-like subfamily of SF2 helicases and interacts with subdomain 1 and 2 to form the RNA binding cleft. NS3h is a multifunctional enzyme, possessing three enzymatic activities: RNA helicase, nucleoside triphosphatase (NTPase), and RNA 5’-triphosphatase (RTPase) (24–29). Previous studies have shown that the two latter activities share a catalytic active site where NTP binds between subdomain 1 and subdomain 2 (26, 30–32). The RNA helicase active site is distinct from that of the NTPase/RTPase active site, located at the helical gate between subdomain 2 and subdomain 3 of NS3 (33–35). The helical gate and β-wedge are responsible for splitting the dsRNA intermediate into two ssRNA strands: the positive-sense viral ssRNA and the negative-sense template ssRNA (33). Single-molecule and structural biology studies have suggested that the negative-sense ssRNA template enters the RNA binding cleft, where the helicase utilizes the energy produced from the hydrolysis of one ATP molecule to power translocation and unwinding of the dsRNA intermediate one base at a time (15, 16, 32, 36–40). A fundamental unanswered question within the SF2 helicase field is how ATP hydrolysis causes structural changes within the helicase to result in translocation and unwinding of dsRNA intermediates.

Between the ATPase active site and the RNA binding cleft, there are eight structural Motifs (I, Ia, II, III, IV, IVa, V and VI) that are highly conserved in the viral/DEAH-like subfamily of SF2 helicases (Figure 1) (22). These structural Motifs play a critical role in substrate binding and are responsible for enzymatic activities within the helicase. One of these Motifs, Motif V, was reported by Davidson *et al.*(24) as a potential link between the ATP binding pocket and the RNA binding cleft due to strong correlations between residues within Motif V and both binding pockets found within the helicase. Additionally, Mastrangelo *et al.*(34) reported that Motif V may be one of the main components of the driving force pulling the helicase along the ssRNA due to conformational changes initiated by the binding of ATP which propagated through Motif V opening the helical gate. At least two residues within Motif V have direct interactions with either ATP hydrolysis active site or the bound ssRNA molecule. G414 coordinates with the lytic water found within the ATPase active site, and T408 interacts with the phosphate backbone of the bound ssRNA (24). In addition to these interactions, Motif V has strong coupling between the other highly conserved Motifs that interact with the ATP binding pocket and the RNA binding cleft suggesting that Motif V may be central to the communication between these two binding pockets (24, 41). Therefore, we investigated the role of Motif V as a critical link between the two binding pockets. We utilized a combination of all-atom molecular dynamics simulations, biochemical assays and virological assays to understand the role of Motif V in internal enzymatic communication leading to helicase function. Our results show residues within Motif V control not only the enzymatic activities of the helicase but also the ability for the flaviviruses to replicate *in vitro*. This is due to importance of the secondary structure of Motif V, as indicated through molecular dynamics simulation.

**Figure 1.**
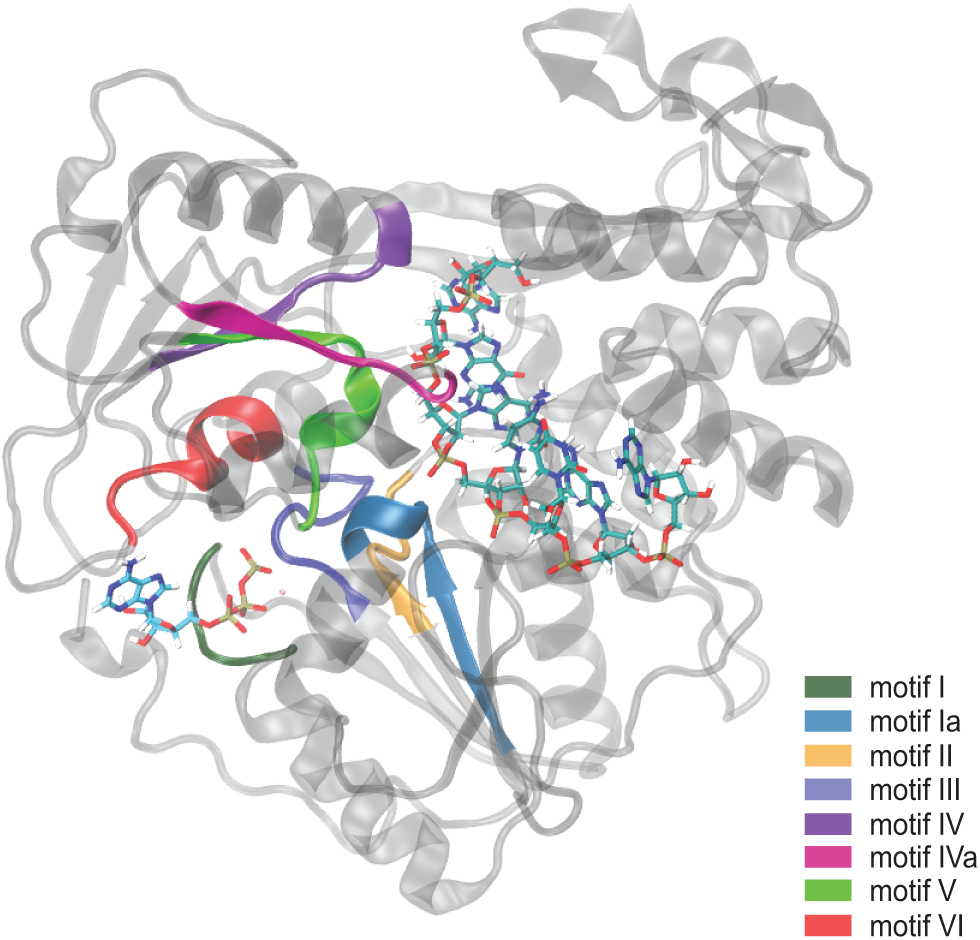
Eight structural motifs of Flavivirus NS3 helicases are highly conserved. The structural Motifs (I, Ia, II, III, IV, IVa, V, VI) are located between the ATP binding pocket and RNA binding cleft. Each Motif is highlighted as follows: Motif I in dark green; Motif Ia in blue; Motif II in orange; Motif III in royal blue; Motif IV in purple; Motif IVa in magenta; Motif V in lime green; Motif VI in red.

## RESULTS

### Motif V Mutants Affect Viral Genome Replication

As previously discussed in Davidson *et al.*, Motif V may play a role in the communication between the ATP binding pocket and the RNA binding cleft/helical gate (24). As one of the highly conserved Motifs in SF2 helicases, Motif V consists of 13 residues spanning position 404 through position 416 (Figure 2A). Eight of these residues are 100% conserved across all flaviviruses, both mosquito and tick-borne (Figure 2B). The other five residues are highly conserved in flaviviruses that infect insects and mammals, and are more variable in tick-borne restricted flaviviruses (Table S1). Interestingly, the conserved residues across all the flaviviruses interact with either the ssRNA or ATP substrates. These highly conserved residues include T408, D409, I410, E412, M413, G414, A415, N416. Of the mostly conserved residues, T407 and S411 mainly interact with each other through a hydrogen bond that is hypothesized to stabilize the α-helical secondary structure of Motif V (Figure 2C). The other mostly conserved residues, F404, V405 and V406, interact primarily with other hydrophobic residues in the surrounding area. Generally, all of Motif V interacts with either a bound substrate or other highly conserved Motifs that interact with bound substrate. Therefore, we investigated the importance of each residue in Motif V on viral replication.

**Figure 2.**
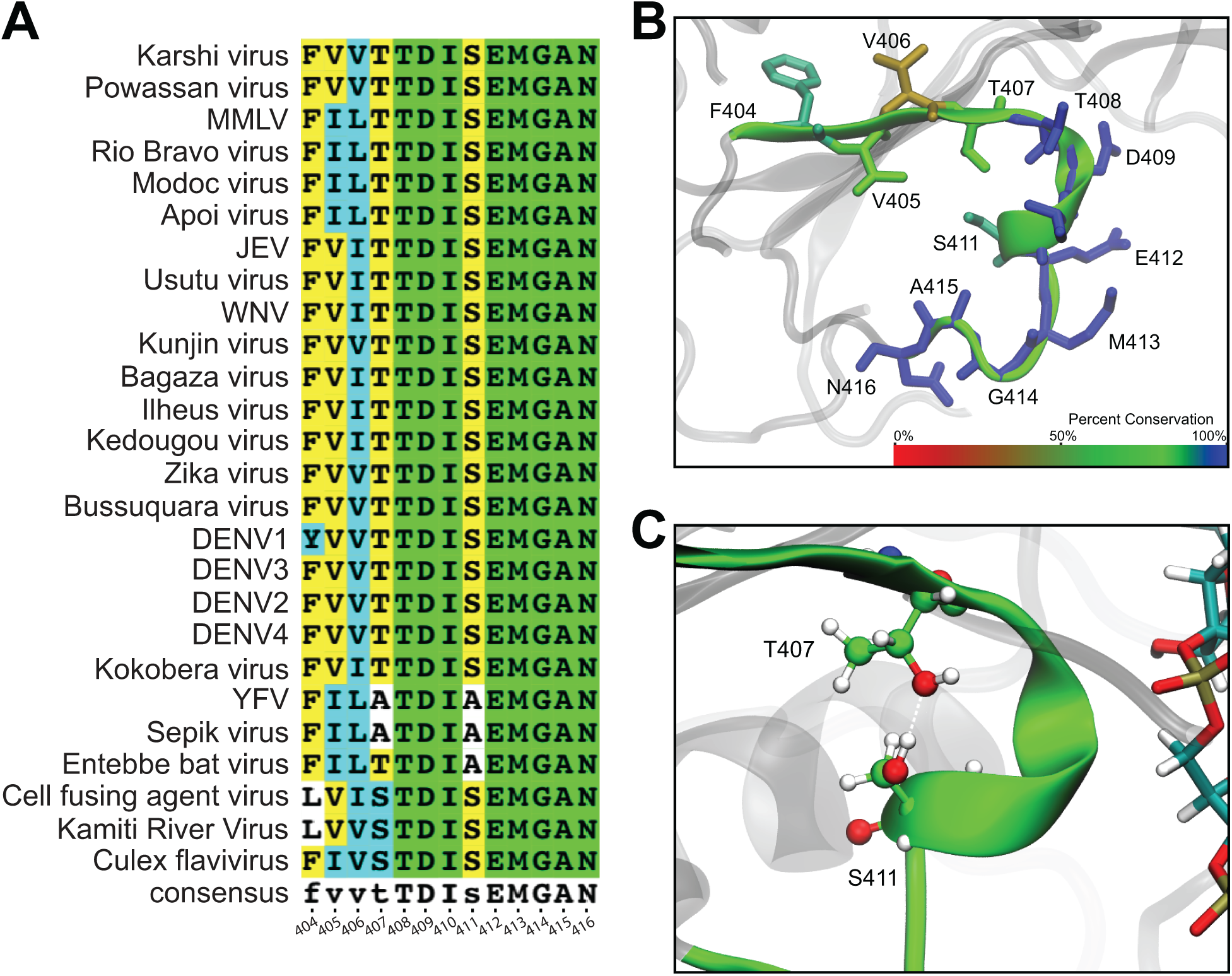
Majority of Motif V residues are highly conserved across all flavivirus NS3 helicases. **A)** The sequence of Motif V from all flaviviruses, multiple-species and insect-restricted. **B)** The percent conservation for each residue within Motif V is projected onto the secondary structure. The color bar ranges linearly from red (0.0% conserved) to blue (100.0% conserved). **C)** T407 and S411 interact through a hydrogen bond.

We individually mutated every residue within NS3h Motif V in a luciferase-expressing WNV replicon system and determined replication efficiency for each mutant. Each Motif V mutation was designed to interrupt specific WT residue-residue interactions or residue-substrate interactions. As a negative control, D664 in the NS5 polymerase catalytic active site was mutated to valine (D664V) in order to disrupt viral genome replication (42–45). Additionally, the NS3 mutants A286L (Motif II) and R387M (Motif IVa) were used as controls for ablating ATPase activity and RNA binding affinity, respectively (30, 46–50). The small hydrophobic sidechain of A286 interacts with other hydrophobic residues stabilizing Motif II interactions with bound ATP. By introducing a bulky hydrophobic residue with A286L, the Motif II interactions with ATP will be perturbed resulting in a disruption in ATP hydrolysis activity. The guanidium head group of R387 interacts with the phosphates of the RNA backbone. By removing the guanidium head group with R387M, important protein-RNA interactions will be interrupted. NS5 D664V and NS3 A286L and R387M ablated viral genome replication, as expected (Figure 3A). All of the Motif V mutants, except for V405M, V406M, T407A, T408S, S411A and A415G, ablated viral genome replication as well. V405M and T407A reduced replication to approximately 2.5%, whereas the I406M and A415G mutants reduce viral genome replication to less than 20%. T408S and S411A were approximately 48% and 39% active, respectively compared to WT NS3h. The results from the viral genome replication were projected onto the structure of Motif V (Figure 3B). We noted that residues with ablated viral replication consisted mostly of the highly conserved residues, suggesting that these highly conserved residues that interact with either the RNA or the ATP play an important role in viral replication. Interestingly, T408 and A415, both highly conserved residues, still maintain a reduced level of viral replication, suggesting that there is not a direct correlation between residue conservation and effect on replication.

**Figure 3.**
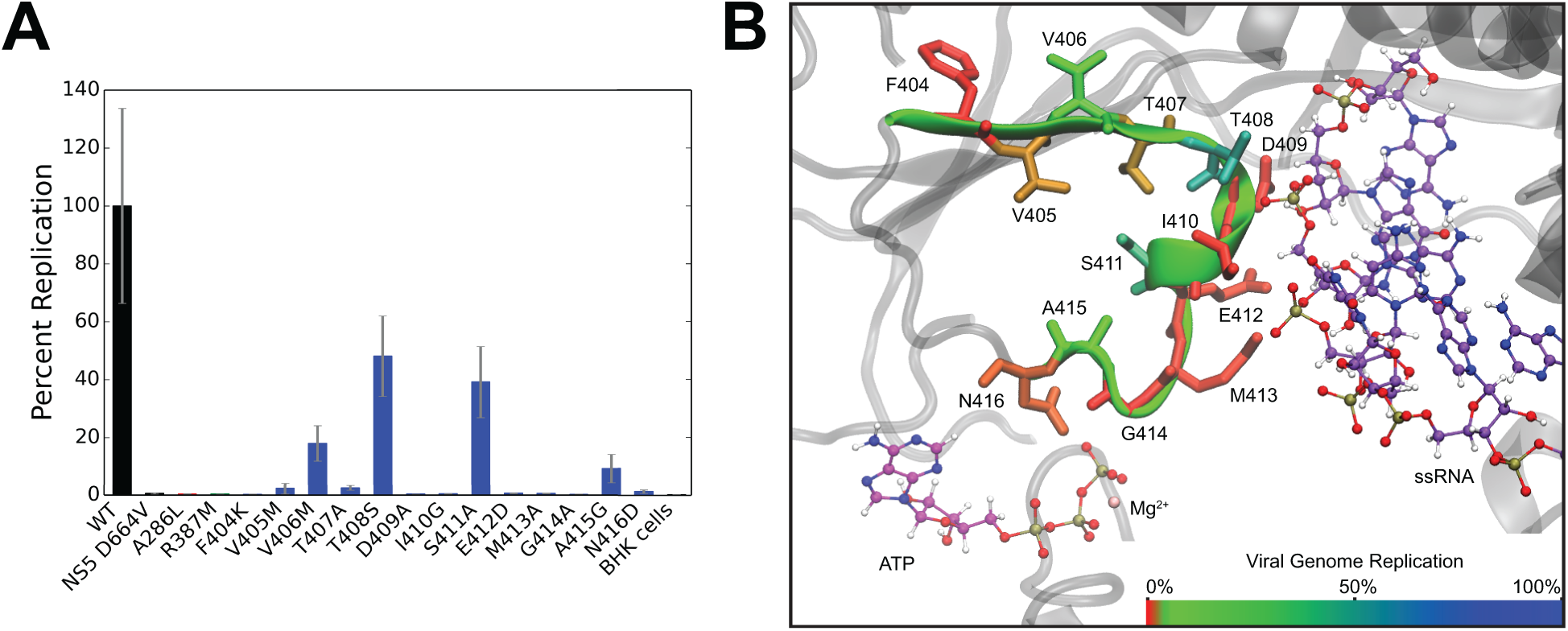
Mutations in Motif V negatively affect viral genome replication. Mutations to remove specific interactions within the central location of Motif V between ATPase active site and RNA binding cleft were tested in **A)** viral genome replication. Results from the viral genome replication were projected onto **B)** the structure of Motif V with the hydrogen bond highlighted between residues T407 and S411. The color bar ranges linearly from red (0% replication) to blue (100% replication). These data suggest that the hydrogen bond between T407 and S411 is important for viral genome replication.

As mentioned before, residues T407 and S411 interact with each other through a hydrogen bond (Figure 2C). However, the T407A mutant nearly ablates genome replication while the S411A mutation only reduces genome replication by approximately 60%. Both mutations should disrupt the hydrogen bond, but T407A and S411A have significantly different replication levels suggesting that these two mutations have other interactions that potentially affect one or more of the NS3h functions during viral replication. These NS3h functions include the ability for the helicase to bind RNA, the ability to hydrolyze ATP, and the ability to unwind the dsRNA intermediate. Therefore, in order to determine whether or not these mutations have an effect on the individual helicase functions, we tested both T407A and S411A biochemically.

### NS3h Residues T407 and S411 Affect ATPase and Helicase Functions but not RNA Binding Affinity

We designed/adapted three biochemical assays to observe how the T407 and S411 mutations would affect the individual helicase functions. Recombinant WT, T407A, S411A, A286L (ATPase control) and R387M (RNA binding control) NS3h were purified and tested in an RNA binding affinity assay, an ATPase activity assay and a helicase unwinding activity assay. We used DENV4 NS3 proteins for this work due to established expression and purification protocols being in place (51) and the high degree of conservation between WNV and DENV in Motif V. To start, the RNA binding affinity of WT, T407A, S411A, A286L, and R387M were tested via a fluorescence polarization assay (52). Fluorescence polarization for a single-stranded RNA oligo labeled with an Alexa-488 fluorophore was measured as the NS3 helicase increased in concentration. The binding affinity or K_d_ for WT NS3h was 1.9 ± 0.8 µM, while the K_d_ for R387M was 76.4 ± 31.0 µM (Table 1A). The R387M K_d_ was significantly different compared to WT, suggesting that R387M does not bind the ssRNA as well as WT. Additionally, a similar trend was observed for WT and A286L. On the other hand, T407A (2.0 ± 1.1 µM) and S411A (1.4 ± 0.2 µM) were not significantly different as compared to WT. This data suggests that neither the T407A and S411A mutations affected ssRNA binding to NS3.

**Table 1.** Enzyme kinetics of WT, A286L, R387M, T407A and S411A NS3 helicase. **A)** The dissociation constant (K_d_) was determined from the RNA binding affinity assay data. **B)** The ATPase activity enzyme kinetics (k_cat_, K_M_, and k_cat_/K_M_) were determined from fitting the data to the substrate inhibition equation. **C)** The helicase unwinding activity enzyme kinetics (k_cat_, K_M_, and k_cat_/K_M_) for each NS3h variant were calculated from fitting the Michaelis-Menten equation to the K_obs_ verse ATP concentration.

| **A** | RNA Binding Affinity |  |  |
| --- | --- | --- | --- |
| | NS3h variant | $K_d$ ($\mu\text{M}$) | Fold Difference |
| | WT | $1.9 \pm 0.8$ | |
| | A286L | $6.7 \pm 0.4$ | 3.5 |
| | R387M | $76.4 \pm 31.0$ | 39.7 |
| | T407A | $2.0 \pm 1.1$ | 1.0 |
| | S411A | $1.4 \pm 0.2$ | 0.7 |

| **B** | ATP Hydrolysis Activity |  |  |  |
| --- | --- | --- | --- | --- |
| | NS3h variant | $k_{\text{cat}}$ ($\text{ms}^{-1}$) | $K_M$ ($\mu\text{M}$) | $k_{\text{cat}}/K_M$ ($\mu\text{M}^{-1}\text{ms}^{-1}$) |
| | WT | $1.2 \pm 0.1$ | $17.3 \pm 4.6$ | $0.0671 \pm 0.0189$ |
| | A286L | $0.9 \pm 0.4$ | $285.1 \pm 150.5$ | $0.0032 \pm 0.0022$ |
| | R387M | $1.6 \pm 0.3$ | $49.5 \pm 21.2$ | $0.0323 \pm 0.0151$ |
| | T407A | $1.2 \pm 0.2$ | $33.7 \pm 12.1$ | $0.0364 \pm 0.0142$ |
| | S411A | $1.8 \pm 0.5$ | $71.5 \pm 34.8$ | $0.0258 \pm 0.0144$ |

| **C** | Helicase Unwinding Activity |  |  |  |
| --- | --- | --- | --- | --- |
| | NS3h variant | $k_{\text{cat}}$ ($\text{ms}^{-1}$) | $K_M$ ($\mu\text{M}$) | $k_{\text{cat}}/K_M$ ($\mu\text{M}^{-1}\text{ms}^{-1}$) |
| | WT | $48.0 \pm 4.0$ | $24.0 \pm 11.0$ | $2.0 \pm 0.8$ |
|  | A286L | 0.0 | 0.0 | 0.0 |
|  | R387M | 0.0 | 0.0 | 0.0 |
| | T407A | $81.0 \pm 6.0$ | $14.0 \pm 6.0$ | $6.0 \pm 2.0$ |
| | S411A | $170.0 \pm 15.0$ | $58.0 \pm 22.0$ | $3.0 \pm 0.9$ |

We next investigated how these mutations affected NS3 ATPase activity. We utilized a colorimetric assay to observe ATP hydrolysis via inorganic phosphate production and release (51). The A286L and R387M mutations were used as controls for this assay as well. The catalytic efficiency (k_cat_/K_M_) for A286L (0.0032 ± 0.0022 µM^−1^ms^−1^) was significantly reduced as compared to WT (0.0671 ± 0.0189 µM^−1^ms^−1^). The twenty-fold decrease in catalytic efficiency was due to a significant increase in the Michaelis-Menten constant (K_M_) for A286L (285.1 ± 150.5 µM) as compared to WT (17.3 ± 4.6 µM) whereas the turnover rate (k_cat_) was consistent between the two systems (Table 1B). This data demonstrates that A286 is critical for ATP hydrolysis. Additionally, the k_cat_/K_M_ for R387M (0.0323 ± 0.0151 µM^−1^ms^−1^) was reduced by a factor of two as compared to WT. Similar to A286L, the slight decrease in the catalytic efficiency of R387M was due to an increase in K_M_ (49.5 ± 21.2 µM), whereas the k_cat_ was not significantly different from WT. Interestingly, the increase in ATPase K_M_ coupled with the decrease in RNA binding K_D_ for R387M as compared to WT suggests that RNA binding promotes ATP binding, consistent with a previous report that ATP hydrolysis is stimulated by RNA (26). When WT NS3 helicase is compared to T407A and S411A, we observe similar turnover rates yet increased Michaelis-Menten constants. These results suggest that T407A and S411A hydrolyze ATP at a similar rate to WT but more substrate is needed for the same catalytic efficiency. Overall, however, ATP hydrolysis was not significantly affected by the T407A or S411A mutations.

Since both RNA binding and ATP hydrolysis were minimally affected by T407A and S411A, we then examined how these mutations affected overall helicase unwinding activity. We utilized a molecular-beacon based helicase activity assay to identify the effects of translocation and unwinding of double-stranded RNA (dsRNA) for WT, A286L, R387M, T407A and S411A (51, 53). WT NS3 helicase unwound dsRNA with a k_cat_ of 48.0 ± 4.0 ms^−1^ and K_M_ of 24.0 ± 11.0 µM (Table 1C). The dsRNA was unwound at a k_cat_ of 81.0 ± 6.0 ms^−1^ and a K_M_ of 14.0 ± 6.0 µM for the T407A mutant, and at a k_cat_ of 170.0 ± 15.0 ms^−1^ and a K_M_ of 58.0 ± 22.0 µM for S411A. These kinetics for T407A and S411A were significantly different compared to WT, which suggests that both mutations are catalytically more efficient than WT NS3h.

### The Interaction Between T407 and S411 Controls NS3 Helicase Function

T407A and S411A exhibit increased helicase activity, so we wanted to investigate if this was due to the absence or presence of a hydrogen bond between the two residues. Therefore, we constructed three additional mutations (T407C, S411C and a double mutant T407C/S411C) in both the replicon and recombinant protein systems to test viral genome replication, RNA binding affinity, ATPase activity, and helicase unwinding activity. Replacing residues T407 and S411 with cysteine residues would largely retain the hydrogen bonding potential for each residue but would remove the methyl group from T407. T407C/S411C can be used to create a disulfide bond (∼ 2 Å) under oxidized conditions, locking 407 and 411 into a closer and more rigid interaction than a hydrogen bond (2.5-3 Å). First, we investigated the effects of T407C, S411C and T407C/S411C on viral genome replication in our replicon system. The viral genome replication of T407C was significantly reduced (13.0 ± 8.0 %), while the double-mutant indicated a reduction to genome replication at 52.7 ± 30.0 % as compared to WT NS3 helicase (Figure 5). On the other hand, S411C exhibited similar genome replication activity as WT. These results suggest that the hydroxyl group of T407 is responsible for part of the effect in genome replication since the cysteine mutation recovers replication by ∼2-3 fold compared to T407A. Additionally, these results indicate that the T407 methyl group may be the important component of the threonine sidechain due to the absence of viral replication in both T407A and T407C, which both lack the methyl group. The S411C mutant shows a complete recovery of replication efficiency, so the effect seen in S411A is due to the removal of the hydroxyl group. This data suggests overall that the hydrogen bond between T407 and S411 plays some role in replication but the methyl in T407 plays a different and potentially more important role in replication. Therefore, we investigated the individual NS3 helicase functions for these mutations in the recombinant protein system to further understand the role of the methyl and hydroxyl groups found in T407 and S411.

**Figure 4.**
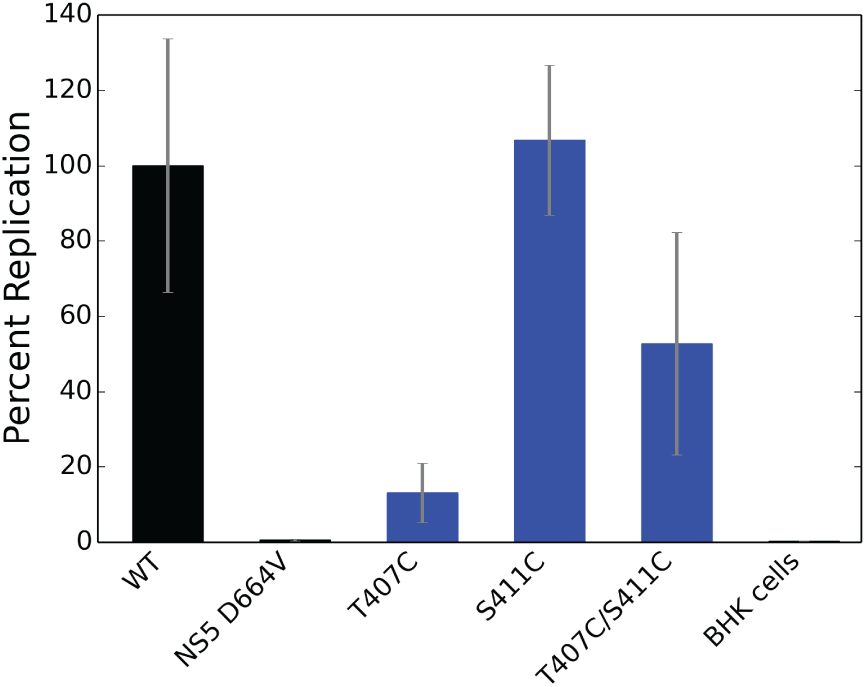
Examining the role of T407 and S411 interaction in NS3h function. Additional mutations to force covalent bonds between the T407 and S411 residues were tested in viral genome replication.

**Figure 5.**
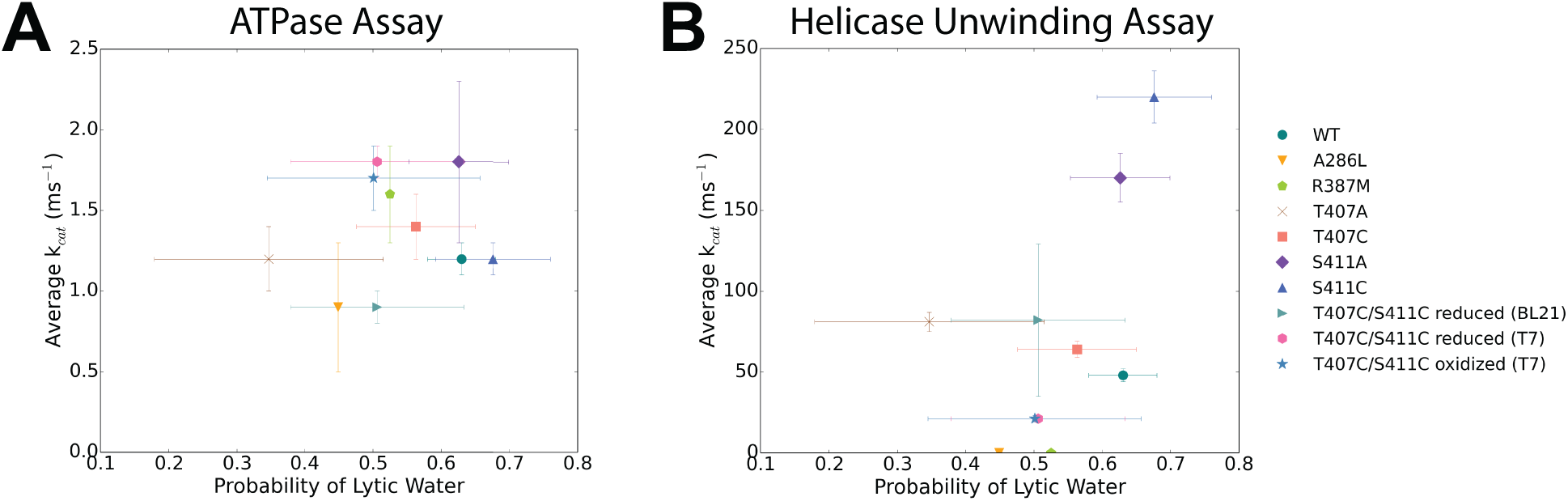
Simulations of mutants do not indicate altered probability of finding a lytic water in the ATPase active site. **A)** The average k_cat_ (ms^−1^) from the ATP hydrolysis activity assay is plotted against finding a lytic water in the ATP binding pocket. **B)** The average k_cat_ (ms^−1^) from the helicase unwinding activity assay is plotted against the probability of finding a lytic water in the ATP binding pocket.

The T407C, S411C, and T407C/S411C mutants were expressed in BL21 DE3 pLysS competent cells to promote a reducing environment. WT and T407C/S411C mutant were also expressed in T7 Shuffle competent cells to promote an oxidizing environment for disulfide bond formation. All of the purified NS3 helicase variants were then tested in the three biochemical assays: RNA binding affinity, ATPase activity and helicase unwinding activity. These cysteine mutations were tested under reducing conditions (TCEP) with the exception of the oxidized WT and T407C/S411C expressed in T7 Shuffle which were tested in both reducing (TCEP) and non-reducing conditions. In all three assays, the BL21 expressed WT compared to T7 Shuffle expressed WT have differing K_d_, k_cat_ and K_M_ values suggesting that the expression cell lines provide a different environment and therefore the T7 Shuffle expressed NS3 helicases cannot be directly compared to the BL21 expressed helicases (Table 2). In the RNA binding affinity assay, the BL21 expressed S411C and T407C/S411C were not significantly different in their binding affinities compared to WT, whereas T407C indicated a weaker RNA binding affinity of 8.0 ± 2.5 µM, which was a 4.2 fold difference compared to WT (Table 2A). Additionally, the reduced T7 expressed T407C/S411C showed a 1.3 fold increase in K_d_ as compared to the reduced T7 Shuffle expressed WT. Similarly, the WT and T407C/S411C expressed in T7 Shuffle cells under non-reducing conditions were also not significantly different with a 0.6 fold difference. All of this data suggest that T407C slightly decreases RNA binding affinity potentially due to the lack of the methyl group, which may play a role in RNA binding to the helicase.

**Table 2.**
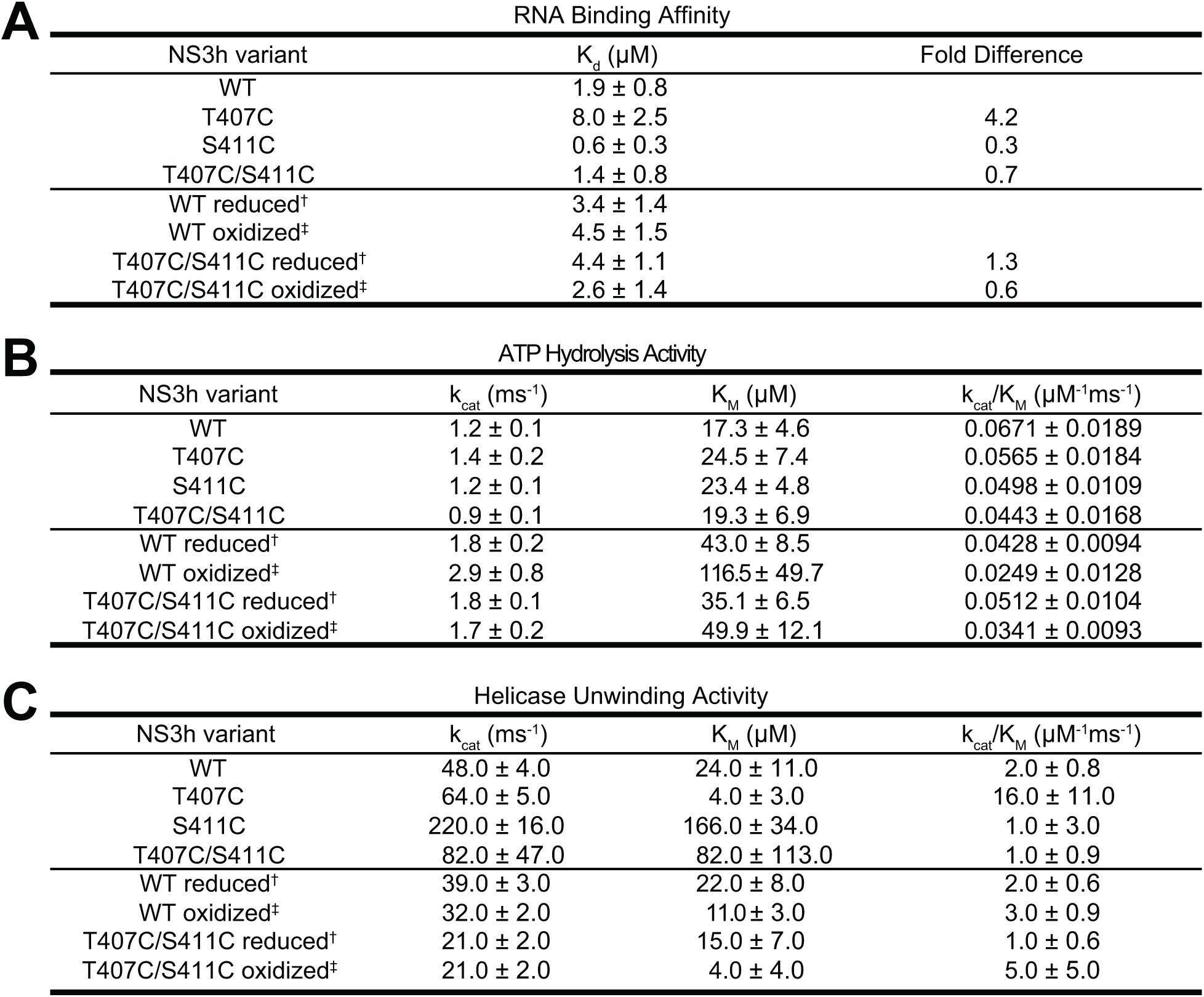
Enzyme kinetics of T407C, S411C, and T407C/S411C NS3 helicase. **A)** The dissociation constant (K_d_) was determined from the RNA binding affinity assay data. **B)** The ATPase activity enzyme kinetics (k_cat_, K_M_, and k_cat_/K_M_) were determined from fitting the data to the substrate inhibition equation. **C)** The helicase unwinding activity enzyme kinetics (k_cat_, K_M_, and k_cat_/K_M_) for each NS3h variant were calculated from fitting the Michaelis-Menten equation to the K_obs_ verses ATP concentration.

| **A** | RNA Binding Affinity |  |  |
| --- | --- | --- | --- |
| | NS3h variant | $K_d$ ($\mu\text{M}$) | Fold Difference |
| | WT | $1.9 \pm 0.8$ | |
| | T407C | $8.0 \pm 2.5$ | 4.2 |
| | S411C | $0.6 \pm 0.3$ | 0.3 |
| | T407C/S411C | $1.4 \pm 0.8$ | 0.7 |
| | WT reduced<sup>†</sup> | $3.4 \pm 1.4$ | |
| | WT oxidized<sup>‡</sup> | $4.5 \pm 1.5$ | |
| | T407C/S411C reduced<sup>†</sup> | $4.4 \pm 1.1$ | 1.3 |
| | T407C/S411C oxidized<sup>‡</sup> | $2.6 \pm 1.4$ | 0.6 |

| **B** | ATP Hydrolysis Activity |  |  |  |
| --- | --- | --- | --- | --- |
| | NS3h variant | $k_{\text{cat}}$ ($\text{ms}^{-1}$) | $K_M$ ($\mu\text{M}$) | $k_{\text{cat}}/K_M$ ($\mu\text{M}^{-1}\text{ms}^{-1}$) |
| | WT | $1.2 \pm 0.1$ | $17.3 \pm 4.6$ | $0.0671 \pm 0.0189$ |
| | T407C | $1.4 \pm 0.2$ | $24.5 \pm 7.4$ | $0.0565 \pm 0.0184$ |
| | S411C | $1.2 \pm 0.1$ | $23.4 \pm 4.8$ | $0.0498 \pm 0.0109$ |
| | T407C/S411C | $0.9 \pm 0.1$ | $19.3 \pm 6.9$ | $0.0443 \pm 0.0168$ |
| | WT reduced<sup>†</sup> | $1.8 \pm 0.2$ | $43.0 \pm 8.5$ | $0.0428 \pm 0.0094$ |
| | WT oxidized<sup>‡</sup> | $2.9 \pm 0.8$ | $116.5 \pm 49.7$ | $0.0249 \pm 0.0128$ |
| | T407C/S411C reduced<sup>†</sup> | $1.8 \pm 0.1$ | $35.1 \pm 6.5$ | $0.0512 \pm 0.0104$ |
| | T407C/S411C oxidized<sup>‡</sup> | $1.7 \pm 0.2$ | $49.9 \pm 12.1$ | $0.0341 \pm 0.0093$ |

| **C** | Helicase Unwinding Activity |  |  |  |
| --- | --- | --- | --- | --- |
| | NS3h variant | $k_{\text{cat}}$ ($\text{ms}^{-1}$) | $K_M$ ($\mu\text{M}$) | $k_{\text{cat}}/K_M$ ($\mu\text{M}^{-1}\text{ms}^{-1}$) |
| | WT | $48.0 \pm 4.0$ | $24.0 \pm 11.0$ | $2.0 \pm 0.8$ |
| | T407C | $64.0 \pm 5.0$ | $4.0 \pm 3.0$ | $16.0 \pm 11.0$ |
| | S411C | $220.0 \pm 16.0$ | $166.0 \pm 34.0$ | $1.0 \pm 3.0$ |
| | T407C/S411C | $82.0 \pm 47.0$ | $82.0 \pm 113.0$ | $1.0 \pm 0.9$ |
| | WT reduced<sup>†</sup> | $39.0 \pm 3.0$ | $22.0 \pm 8.0$ | $2.0 \pm 0.6$ |
| | WT oxidized<sup>‡</sup> | $32.0 \pm 2.0$ | $11.0 \pm 3.0$ | $3.0 \pm 0.9$ |
| | T407C/S411C reduced<sup>†</sup> | $21.0 \pm 2.0$ | $15.0 \pm 7.0$ | $1.0 \pm 0.6$ |
| | T407C/S411C oxidized<sup>‡</sup> | $21.0 \pm 2.0$ | $4.0 \pm 4.0$ | $5.0 \pm 5.0$ |

The NS3 helicase mutants expressed in BL21 and T7 Shuffle competent cells were tested for their ability to hydrolyze ATP. T7 Shuffle expressed NS3 helicases were tested in both oxidizing and reducing conditions. T407C, S411C, and T407C/S411C expressed in BL21 cells all showed insignificant differences in both k_cat_ and K_M_ as compared to WT. The k_cat_ for oxidized WT and T407C/S411C were not significantly different with a k_cat_ of 2.9 ± 0.8 ms^−1^ and a k_cat_ of 1.7 ± 0.2 ms^−1^, respectively, whereas the K_M_ for WT (116.5 ± 49.7 µM) and T407C/S411C (49.9 ± 12.1 µM) under oxidized conditions were significantly different (Table 2B), suggesting that the disulfide bond present in the oxidized form of T407C/S411C increases ATP hydrolysis activity. On the other hand, WT and T407C/S411C expressed in T7 Shuffle cells under reduced conditions exhibit an insignificant difference for either k_cat_ and K_M_. Overall, these data suggest that hydrolysis of ATP is similar between WT and the cysteine mutations meaning that the cysteine residues do not influence ATP hydrolysis directly.

Finally, the various cysteine NS3 helicase mutants as well as WT expressed in both BL21 and T7 Shuffle cells were tested in the helicase unwinding activity assay. When comparing T407C, S411C and T407C/S411C expressed in BL21 cells to WT, we observe a higher k_cat_ with a lower K_M_ for each mutant except for T407C, which exhibits a higher K_M_ value (Table 2C). This data suggest that both S411C and T407C/S411C unwind dsRNA faster than WT even though they do not bind dsRNA as well. T407C also unwinds dsRNA faster than WT, but the affinity for RNA is stronger suggesting that T407C is a more catalytically efficient helicase overall compared to WT. The catalytic efficiency of WT and T407C/S411C expressed in T7 cells indicates no significant difference in the ability to unwind dsRNA substrates suggesting that the double-mutant exhibits similar overall unwinding activity compared to WT.

### In silico mutations of T407 and S411 validate experimental assays

We further investigated how each mutant, alanine and cysteine, affected either the binding energy of the substrates or the overall secondary structure of NS3 helicase in the ssRNA+ATP substrate state using all-atom molecular dynamics simulations. Each mutant (A286L, R387M, T407A, T407C, S411A, S411C, T407C/S411C) was simulated in triplicate for 1 μs using the ff14SB force field within the AMBER18 software package. Once the simulations were completed, the root-mean squared deviation (RMSD) was calculated in reference to the initial starting structure to determine the equilibration time that would be excluded for all other analyses. The three analyses performed on the simulations were a nonbonding interaction energy analysis, a probability of finding a lytic water in the ATP binding pocket, and a projected covariance magnitude analysis. These analyses were specifically designed to be compared results between simulation and the three biochemical assays.

First, we utilized the nonbonding interaction energy analysis to provide insight into how the mutations affect RNA binding affinity from simulation. The nonbonding interaction energy was calculated between the bound ssRNA and the entire protein for each NS3 helicase variant. The more negative the linear interaction energy, the more strongly bound the ssRNA substrate is in the helicase RNA binding cleft. All of the mutations exhibit strong ssRNA binding as compared to WT with the exception of A286L, R387M and T407C mutants (Table 3A). The A286L and R387M mutations increase in their nonbonding interaction energies as compared to WT by 5.3% and 13.8%, respectively. These data suggest that A286L and R387M bind ssRNA weakly compared to WT. Additionally, T407C exhibits a nonbonding interaction energy of - 708.6 ± 36.7 kcal/mol. The more positive nonbonding interaction energy suggests that T407C binds ssRNA more weakly than WT. The lower nonbonding interaction energy correlates well with the reduced RNA binding affinity for T407C (Table 2A), validating our computational approach. Additionally, we observed an increase in sidechain mobility in the T407C mutant simulation compared to WT (Figure S1), suggesting that the threonine at position 407 may control fluctuations within Motif V allowing for the helicase to bind ssRNA strongly.

**Table 3.**
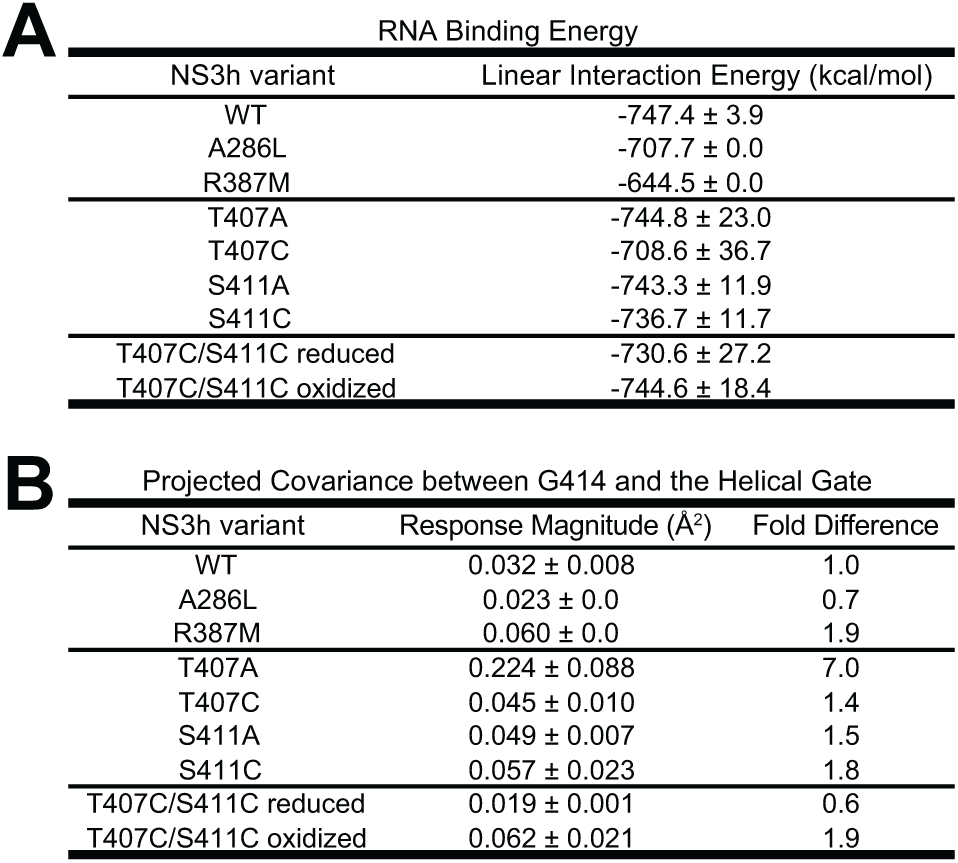
T407 and S411 mutations do not affect nonbonding interaction energies of bound ssRNA to NS3 helicase. **A)** The nonbonding interaction energies were calculated between all residue in the protein to all nucleotides of the single-stranded RNA substrate bound into the RNA binding cleft of NS3 helicase. The post-analysis AMBER18 Tools package, cpptraj, was utilized in order to calculate the linear interaction energy (kcal/mol) for each mutation run in triplicate with an interaction cutoff of 12 Å and short-range, electrostatic energies were calculated with a dielectric of 1. The resulting linear interaction energies were average over the three simulations. **B)** The projected covariance magnitude represents the fluctuations of G414 and residues S364 and K366 projected onto the helical gate access site axis. Each mutant is reported here as well as the fold-difference compared to WT NS3h.

Next, we investigated how the mutations affect the ATP binding pocket through determining the probability of finding a lytic water in the ATP hydrolysis active site. In order to do this, we utilized three collective variables to select for lytic waters: (1) the nucleophilic attack distance between the water oxygen atom and the ATP-γ-phosphate atom, (2) the nucleophilic attack angle between the water oxygen atom and the terminal phosphoanhydride bond of ATP, and (3) the angle between the water dipole moment and the terminal phosphoanhydride bond of ATP. Once the lytic waters were determined, the simulations were analyzed to determine the probability of finding a lytic water throughout the entire equilibrated portion of the simulations. The WT simulations indicate that 63 ± 5% of the time, a lytic water was found within the ATP binding pocket. All of the mutations are not significantly different compared to the WT, except for A286L (44.9 ± 0.11%), R387M (52.5 ± 0.12%), and T407A (34.7 ± 16.8%). These probabilities suggest that A286L, R387M, and T407A mutations have negatively affected the ATP hydrolysis active site due to the low probability of finding a lytic water in the ATP binding pocket during the simulations. However, when we compare the probability of finding a lytic water to the turnover rate determined from the ATPase assay, we observe no significant difference between the mutations and the WT NS3 helicase (Figure 5A). On the other hand, comparing the probability of finding the lytic water to the turnover rate determined from the helicase assay, we observe S411A and S411C are set apart from the rest of the mutations, suggesting that these two mutations are catalytically more efficient than WT and the other mutations because they are more likely to have a lytic water in the ATPase active site (Figure 5B).

Lastly, we implemented a projected covariance analysis to examine how the mutants affect the dynamics of the helical gate through changes in fluctuations between the ATPase active site and the helical gate. As previously mentioned, during replication, the helical gate and β-wedge are responsible for unwinding the dsRNA intermediate into two ssRNA molecules (33). One of the two strands enters into the RNA binding cleft through the helical gate access site. The ssRNA strand in the access site interacts with surrounding residues S364, I365, K366, D603, P604, and M605 found within two α-helices (α-helix 2 in subdomain 2 (α2’) and α-helix 6 in subdomain 3 (α6”)) that flank either side of the incoming ssRNA (34). Previous studies have suggested that the dynamics of the helical gate are controlled by the presence of bound ATP (34). Therefore, we wanted to determine how the mutations within Motif V affect the connection between the ATP binding pocket and the helical gate. We quantified this connection as the positional covariance of ATP pocket residues (sources) and helical gate residues (sinks). Specifically, we investigated the covariance of G414 (Motif V, source) and S364 and K366 (α2’, sink residues) along the helical gate opening/closing coordinate. This quantified as the average of the projections of the covariance tensors between source and sinks along helical gate opening/closing direction (Table 3B).

The WT simulations indicated a projected covariance of 0.032 ± 0.008 Å^2^ suggesting that there was a correlation between the fluctuations seen between the ATP binding pocket and the helical gate. Both T407C (0.045 ± 0.010 Å^2^) and S411C (0.057 ± 0.023 Å^2^) were not significantly different compared to WT, even though their fold differences indicated an increase in projected covariance along the helical access. All of the rest of the mutations were significantly different compared to WT (Table 3B). Both A286L (0.023 ± 0.0 Å^2^) and reduced T407C/S411C (0.019 ± 0.001 Å^2^) showed a decrease in fold difference (0.7 and 0.6 respectively) compared to WT, suggesting that these mutations negatively affect the dynamics of the helical gate (Table 3B). R387M, T407A, S411A and oxidized T407C/S411C indicate an increase in fold difference (1.9, 7.0, 1.5 and 1.9 respectively) compared to WT (Table 3B), suggesting that the fluctuations through Motif V positively affect the motion of the helical gate promoting translocation and unwinding of the dsRNA intermediate. Interestingly, when we compared the projected covariance magnitude fold differences to the turnover rate fold differences of the helicase unwinding activity assay, all of the data support each other except for A286L, R387M and oxidized T407C/S411C. Both A286L and R387M were unable to unwind dsRNA in the helicase unwinding assay, but in simulation these mutants indicated a decrease and an increase in the projected covariance along the helical gate axis, respectively. The difference observed between simulation and experiment may be due to the pre-bound state of the simulation, whereas in experiment the helicase still needs to bind the RNA. Additionally, the turnover rate of the oxidized double mutant indicated a decreased rate compared to WT, while the projected covariance magnitude indicated an increased fluctuation projected onto the helical axis compared to WT. This discrepancy might be due to the inability to control the state of the disulfide bond in experiment.

## DISCUSSION

In this study, a combination of computational, biochemical and virological experiments were utilized to investigate the role of Motif V in NS3 helicase functions. WT simulations pointed to Motif V as a key player in the communication between the ATP binding pocket and the RNA binding cleft. To determine how Motif V may play a role in the communication, mutations were introduced in both replicon and recombinant protein systems. The mutations were tested in a viral genome replication assay and results suggested that the residues within Motif V affect how the virus replicates. From the viral replication assay, residues T407 and S411 sparked our interest due to the differing replication activity with prior knowledge of their interaction through a hydrogen bond that may stabilize Motif V structure. We tested both alanine and cysteine mutations in an RNA binding affinity assay, an ATP hydrolysis activity assay and a helicase unwinding activity assay as well as simulated theses mutations in all-atom molecular dynamics. Results in this paper suggest that the methyl group in the sidechain of T407 may play a role in decreasing helicase unwinding activity through stabilizing interactions with residues T408, K366, and R387. These residues all interact with the bound RNA allowing for viral replication. We can directly compare between the helicase unwinding assay and viral replication assay because the helicase rates are physiologically relevant given the viral replication rate. Overall, our data suggests that Motif V is critical for the communication between the ATP binding pocket and the RNA binding cleft in NS3 helicase.

Initial analyses of WT simulations supported results reported in Davidson *et al.* that Motif V may play a critical role in the communication between the ATP binding pocket and the RNA binding cleft (24). This led us to mutating all of Motif V residues (404 to 416) in the replicon system. Interestingly, only two Motif V mutations have been studied previously within the *Flaviviridae* virus family: T411A in HCV and G414A in dengue 2 virus (30, 54). Both of these mutations were tested biochemically but were not tested for their effect on viral genome replication. Therefore, our replicon data represents the first replication-based examination of the effect of Motif V mutations on viral genome replication. We observed that the majority of the mutations ablated viral replication activity with the exception of residues V405, V406, T407, T408, S411 and A415. Interestingly, most of the highly conserved residues ablated viral replication, but not all did, suggesting that the highly conserved residues can have some variability with the type of amino acid at those positions. Of the residues that did not ablate activity, T407 and S411 were of most interest due to the variable nature of these residues across all flaviviruses and the hydrogen bond that may stabilize the secondary structure of Motif V during replication. Therefore, we focused on these two positions for the remainder of the discussion.

T407 and S411 were mutated to alanine residues in the recombinant NS3 helicase and were expressed, purified and tested in RNA binding affinity, ATPase activity, and helicase unwinding activity assays along with two controls A286L and R387M. These three assays provide specific insight into how the helicase binds ssRNA and the rates of (1) ATP hydrolysis and (2) dsRNA unwinding. The rates determined from the latter two assays cannot be directly compared because the time scales are different due to the fact that the ATPase assay is an end-point assay while the helicase unwinding activity assay is a continuous assay. The results from the RNA binding affinity assay indicated that T407A and S411A bind ssRNA as strongly as WT. The results from the ATP hydrolysis activity assay indicated that T407A and S411A hydrolyze ATP as well as WT. On the other hand, T407A and S411A seem to unwind dsRNA more quickly than WT in the molecular beacon helicase assay, suggesting that T407 and S411 are involved in the linkage between the ATP binding pocket and the RNA binding cleft. If the individual activities of RNA binding and ATP hydrolysis are unaffected by these mutations but the over ability for the helicase to unwind dsRNA is affected, then these two mutations must play a role in the communication between the substrate binding pockets. The observation that T407 and S411 increased helicase turnover rate was surprising and suggested that one or both of these residues may naturally act to weaken the effect of ATP hydrolysis on helicase activity giving raise to the intriguing possibility that the helicase has evolved to slow down its ATPase-dependent helicase activity. Position 407 is found predominately as a threonine and 411 found as serine (Figure 3A and Table S1), but in a small group of flaviviruses position 407 can be alanine or serine and 411 can be alanine. This may indicate that the helicase function in other flaviviruses (yellow fever virus, Sepik virus, Entebbe Bat virus, Cell Fusing Agent, Kamiti River virus, and Culex flavivirus) could be faster than most other flaviviruses.

We further investigated which aspects of the threonine and serine residues may play a critical role in the regulation of helicase unwinding activity by introducing cysteines at positions 407 and 411 individually or as an double-mutant, T407C/S411C. T407C, S411C and T407C/S411C were mutated in both the replicon and recombinant protein systems. We observed an increase in replication activity for the T407C mutation, a full recovery for the S411C mutation, and an increase in replication activity for the double-mutant. T407C was the only mutation that weakens the ability of the helicase to bind ssRNA. The ATP hydrolysis activity is unchanged for all of the cysteine mutations, and the helicase unwinding activity indicate T407C and S411C increase unwinding activity while the double mutant under oxidizing and reducing conditions remain as active as WT. The combination of these data suggest that the methyl group in the sidechain of T407 may be important for slowing down helicase function. We observed a catalytically more efficient T407A in unwinding dsRNA, but a significant reduction in viral replication activity. When we mutate T407 to a cysteine (similar to a threonine but lacking in the methyl group), we recover some replication activity. When we mutated S411 to cysteine, we completely recover viral replication suggesting that while the hydrogen bond between T407 and S411 is important, T407 has a separate function related to its methyl group. The methyl group in the T407 sidechain normally interacts with the phenylalanine ring of F361 via hydrophobic interactions, potentially stabilizing the residues T408, K366, and R387, which all interact with the RNA phosphate backbone (Figure S1A). By removing the methyl from T407 with the T407C mutation, the sidechain of position 407 is able to rotate and interact not only with F361 but also L385, potentially causing RNA interacting residues to become more variable with their interactions with bound ssRNA. As a result, these three residues that interact with the RNA may become more catalytically efficient in unwinding the dsRNA substrate. In the T407C mutation, the viral replication recovers some activity suggesting that the hydroxyl group in T407 is also potentially important for proper viral replication due to the ability to form a hydrogen bond like interaction with S411. Overall, when the helicase becomes catalytically more efficient in the absence of the T407 methyl, viral genome replication decreases. It is unclear why this is the case, but one hypothesis is that reduced replication may be due to a rate mismatch between T407A, T407C, and S411A NS3 and the RNA-dependent RNA polymerase of NS5. If this is the case, it may suggest that the NS3 helicase and NS5 polymerase have co-evolved to match their catalytic rates so that RNA is fed at an optimal rate between the two enzymes.

All of the NS3 helicase mutations were simulated using the ff14SB force field within the AMBER18 software package to better define how the mutations affected NS3 helicase function. Three analyses were performed on the simulations in order to determine how the structure of the helicase has changed due to the presence of the mutations. The first analysis we utilized was the nonbonding interaction energy analysis. The interaction energies determined from this analysis were compared to the RNA binding affinity (K_d_) obtained from experiment. We observed that in both computation and experiment, the mutations (T407A, T407C, S411A, S411C, and T407C/S411C) did not significantly change the ability for the helicase to bind ssRNA. The only mutation that weakened the binding of ssRNA in the helicase was R387M, which was expected because this mutation was a control for RNA binding. As we moved forward with our analysis of the simulations, we investigated how the mutations affected the probability of finding a lytic water in the ATPase active site (Figure 5). When we compared the probability of finding a lytic water in the ATP binding pocket to the turnover rate determined from the ATPase assay, we observed that none of the mutations significantly changed the probability of finding a lytic water. When comparing the probability of a lytic water to the turnover rate of the helicase activity, we observed that S411A and S411C were set apart from the rest of the mutations suggesting that these two mutations play a role in controlling the rate of unwinding activity through the hydrogen bond with T407. Interestingly, when T407 is mutated to a cysteine, we observe an increase in mobility at position 407 (Figure S1). The cysteine residue lacks the methyl group and contains a thiol group instead of a hydroxyl group found in the threonine sidechain. The lack of the methyl group seems to allow for the thiol group to fluctuate frequently throughout the simulation (Figure S1B and S1C), suggesting a potential hydrophobic interaction between T407 and nearby hydrophobic residues (such as L385) that may influence the position of T407 within Motif V. This supports our previous results for T407 in that the methyl groups orientation is critical for helicase function. Additionally, the projected covariance magnitude analysis indicated that T407A increased by 7.0 compared to WT suggesting that the methyl and hydroxyl group of threonine are important for stabilizing the interactions of position 407 with its surrounding residues. Without either of those functional groups at position 407, Motif V may allow more structural changes to occur that positively affect the helical gate. This is evident when we take into account the projected covariance of T407C. T407C indicated a slight increase compared to WT, which is similar to what is observed in the k_cat_ of T407C from the helicase unwinding assay. The thiol group of the cysteine potentially interacts with S411 stabilizing Motif V enough to control the structural changes within the helicase to allow for optimal translocation and unwinding of dsRNA.

This work provides detailed computational, biochemical, and virological insight into how the helicase utilizes the energy produced from ATP hydrolysis to power translocation and unwinding of the viral double-stranded RNA intermediates. We have learned that the methyl group in T407 is potentially critical for regulating the helicase to maintain an optimal rate of translocation and unwinding during replication. If the T407 is mutated to an alanine or a cysteine, the viruses we examined are not able to replicate the viral genome efficiently *in vitro*. This T407 residue within Motif V interacts with S411, which seems to properly orient T407 potentially stabilizing the structure of Motif V. When we remove that interaction between T407 and S411 with a mutation to S411, the helicase is able to unwind dsRNA intermediates with more catalytic efficiency suggesting that T407 is responsible for regulating translocation and unwinding dsRNA intermediates. As previously mentioned, the role of Motif V in SF2 helicases is not well understood. This work has provided computationally driven experimental insight into how Motif V plays a role in the communication between the ATP binding pocket and the RNA binding cleft in the viral-DEAH subfamily of SF2 helicases (22). Additional work will be required to determine if the results we observe here for flaviviruses hold in other subfamilies of SF2 helicases. Ultimately, the data from this paper suggests that flaviviruses may utilize suboptimal NS3 helicase activity for optimal genome replication, and that Motif V may play an inhibitory role for helicase activity during viral infection *in vitro* and *in vivo*. The role of Motif V in controlling helicase function will need to be further defined, which may eventually lead to the development of novel vaccine candidates or antiviral drug targets.

## EXPERIMENTAL PROCEDURES

### Plasmids and Mutagenesis

The DNA expression plasmid used in the protein purification protocol for dengue 4 NS3 (Accession Number: AQV12689) was adapted from Luo and co-workers (6). The QuikChange mutagenesis (Agilent Technologies) method was used to mutate specific residues (A286L, R387M, T407A, T407C, S411A, S411C, and T407C/S411C) within the WT dengue NS3h plasmid optimized for expression in *E. coli*. Plasmids for each mutation were submitted to GENEWIZ for sequencing to verify each sequence.

### Protein Expression and Purification

WT dengue NS3h gene with an N-terminal thioredoxin, 6X-histidine tag, thrombin cleavage site was synthesized and cloned into a T7 expression plasmid. The wild-type (WT) NS3h construct was transformed into BL21 DE3 pLysS or T7 Shuffle *E. coli* competent cells. 5 mL Luria broth (LB) cultures containing ampicillin (50 μg/mL) and chloramphenicol (34 μg/mL) were grown overnight at 37 °C, and the next day the entire 5 ml culture was added to a 750 mL LB culture containing 50 μg/mL of ampicillin. 750 mL cultures were incubated until an OD_600_ of 0.6 was reached, the cultures were induced with a final concentration of 400 µM IPTG for 18 hours, and bacteria collected by centrifugation. Bacteria were resuspended with 20 mL low imidazole buffer (LIB) (50 mM Tris-Base pH 8.00, 400 mM NaCl, 10 mM imidazole, 5% glycerol, and 12 mM CaCl_2_ and stored at −80°C until use. Pre and post-induction samples were collected and used to verify protein expression by SDS-PAGE gel analysis.

For the protein purification, LIB (20 mL) was added to the frozen bacterial pellets. Once thawed, bacterial cells were disrupted 3 times on a 110S Microfluidizer (Microfluidics), the lysate clarified by centrifugation using a JA-25.50 rotor for 20 minutes at 17,000 rpm at 4 °C, and supernatant filtered using a 0.45 μM syringe filter. Protein purification was achieved using a multi-step approach. The thioredoxin-dengue NS3h complex was first purified from the filtered lysate with nickel affinity chromatography using a HisTrap HP (GE) column on an AKTA Pure FPLC system (GE). The protein complex was eluted off the Ni-column with a linear gradient of high imidazole buffer (HIB) (50 mM Tris-Base pH 8.00, 400 mM NaCl, 10 mM imidazole, and 5% glycerol). Fractions containing the NS3h complex were collected, dialyzed against 350 ml of LIB containing CaCl_2_ for two hours using a Slide-A-Lyzer Dialysis Cassette (3-12 mL (Life Tech)). 75 μL of soluble thrombin (GE Life Sciences) was added to the dialyzed protein for an overnight incubation at 4°C. A second Ni-column was subsequently used to remove thioredoxin from NS3h, which was collected in the flow-through fraction. The flow through fraction was concentrated using a Vivaspin Turbo 15 ultrafiltration spin column (Satorius), then applied to a HiLoad 16/600 Superdex 200 pg (GE Life Sciences) gel filtration column to further purify and buffer exchange the isolated NS3h in gel filtration buffer (50 mM Hepes at pH 8.00, 400 mM NaCl and 20% glycerol). Purified NS3h was concentrated in Vivaspin spin columns before storage at −80°C in single use aliquots. All buffers were made with RNAse free H_2_O. The concentration of NS3h variants were confirmed on by SDS-PAGE (Figure S2A). The recombinant WT NS3h was also confirmed by mass spectrometry (data not shown).

### RNA Helicase Unwinding Activity Assay

For the ATP dependent helicase assay(51, 53), all reactions were performed in 96-well black microplates (Thermo Scientific) in a final volume of 120 μL and they contained 50 nM NS3h, 0.05 mM TCEP, 0.01% Tween20, 5 μg/μL BSA, 1.25 mM MgCl_2_, 25 mM MOPS (pH 6.5), 5 nM RNA complex, and varying concentrations of ATP (Jena Bioscience) ranging from 1 µM to 1 mM. The RNA complex consists of an RNA tagged Alexa-488 oligo (5’-AGUGCGCU-GUAUCGUCAAGG-CACU-3’-AlexF488N) and a DNA tagged Iowa Black Fluorescent Quencher (5’-IABkFQ-CCTACGCCACCAGCTCCGTAGG-3’) bound to an RNA template (5’-GGAGCUGGUGGC-GUAGGCAAGAGUGCCUUGACGAUACAGCUUUUUUUUUUUUUUUUUUUU-3’). The RNA complex was annealed through a slow cooling process by boiling H_2_O to 95°C and allowing to cool to room temperature over an hour. The control with no MgCl_2_ contained 20 mM EDTA. Reactions were scanned individually at 37°C using a Victor X5 multilabel plate reader (Perkin Elmer) instrument for 300 seconds (Figure S2B). Rates were determined from where the progress curves were exponential and linear with time. Data were then fit to the Michaelis-Menten equation (eq 1) using Python2.7

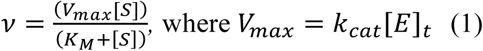

and the *ν* is the velocity, k_cat_ is the apparent first-order rate constant in s^−1^, [E]_t_ is the concentration of the enzyme, [S] is the concentration of the substrate, K_M_ is the concentration of the substrate at one-half k_cat_. The Michaelis-Menten constant (K_M_), turnover rate (k_cat_), and the specificity constant (k_cat_/K_M_) were determined from fitting to equation 1.

### RNA ATPase Activity Assay

A colorimetric ATPase activity assay was derived from the RNA helicase assay (51). The contents of the reactions remained the same. Using a 96-well clear plate, reactions were injected with BIOMOL Green (Enzo), incubated at 37°C for 15 seconds before measuring the absorbance at 650 nm on a Victor X5 multilabel plate reader. To obtain an entire curve for an individual ATP concentration, reactions were measured every 90 seconds for 12.5 minutes (Figure S2C). Rates were determined from progress curves. Data were then fit to the substrate inhibition equation (eq 2) using the R software environment (55)

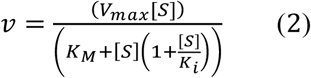

where K_i_ is the inhibition constant. From the fit, the k_cat_, K_M_, k_cat_/K_M_ and the K_i_ were determined for the ATPase assay. K_i_ data is reported in Table S2.

### RNA Binding Affinity Assay

A fluorescence polarization (FP) assay was also adapted from the RNA helicase assay to determine RNA binding K_d_ as previously described (52). The FP assay reactions contained 50 nM NS3h, 0.05 mM TCEP, 0.01% Tween20, 5 μg/μL BSA, 1.25 mM MgCl_2_, 25 mM MOPS (pH 6.5) and 5 nM ppAGUAA tagged Alexa-488 RNA oligo. All reactions were incubated at 37°C for 10 minutes in a 384-well black plate within the plate reader before measuring the fluorescence polarization of the Alexa-488 fluorophore (Figure S2D). Data was analyzed using nonlinear regression analysis in Prism8.

### All-atom Molecular Dynamics Simulations

An exemplar structure of the simulated DENV4 NS3h ssRNA+ATP substrate state (2JLV) was used for all simulations (24, 33). The ssRNA+ATP state was run for 9 systems of NS3h: WT, A286L, R387M, T407A, T407C, S411A, S411C and the double mutant T407C/S411C in oxidized and reduced conditions in triplicate for 1 μs each. All of the mutations were generated from the WT NS3 helicase.

All of the NS3h systems were simulated in explicit solvent MD using AMBER18 software package: ff14SB (protein) and RNA.OL3 (RNA) (56). Parameterization files for the ATP molecule (57) were obtained from the AMBER parameter database. Each system was solvated with a cubic box of TIP3P water with an average dimension of 85 Å. Sodium and chloride ions were added at a concentration of 100 mM to neutralize the system. The simulations were performed with periodic boundary conditions in an isothermal-isobaric (NPT) ensemble with a stochastic barostat of 1 bar and a Langevin thermostat of 310.0 K. The nonbonding interactions cutoff was 12 Å; particle mesh Ewald was used for treating long range electrostatics; and hydrogens were constrained with the SHAKE algorithm. An integration timestep of 2 fs was used. Positions and energies were written every 2 ps.

All systems were initially minimized with a 10,000 step minimization with harmonic restraints (force constant of 75 kcal mol^−1^Å^−2^) placed on all solute atoms and a 10,000 step minimization with no restraints. Following the minimization, each system was heated from 50.0 K to 310.0 K in 10 K increments with a harmonic restraint of 75 kcal mol^−1^Å^−2^ on all solute atoms. Lastly, a series of five equilibration steps were implemented to slowly remove the harmonic restraint.

### Data Analysis of MD simulations

All analyses of MD simulations were performed using Python 2.7 and the MDAnalysis package (58, 59). These analyses include the linear interaction energy analysis, the lytic water analysis and the projected covariance magnitude analysis. Further details of each analysis, except for the linear interaction energy, were described below. Plots were created using Matplotlib(60), and VMD(61) was utilized to visualize simulations and generate structural images. All scripts for the analyses are available on GitHub (https://github.com/mccullaghlab/T407_S411_Mutants_of_NS3h).

Waters within the NTPase active site were defined as “lytic” using three collective variables: (1) the nucleophilic attack distance between the water oxygen atom and the ATP γ-phosphorous atom, (2) the nucleophilic attack angle between the water oxygen atom and terminal phosphoanhydride bond of ATP, and (3) the dipole moment angle between the water molecule’s dipole moment vector and the terminal phosphoanhydride bond of ATP. The first two metrics describe the geometric positioning of waters within the hydrolysis active site. Waters that have a nucleophilic attack distance less than 5Å and a nucleophilic attack angle greater than 160 degrees have the potential to be the lytic water due to their positioning relative to the terminal phosphate group. The third metric describes the chemically relevant orientation of a water in regard to its nucleophilic attack on this phosphate group. An ideally positioned water with a dipole moment angle of greater than 90 degrees is defined as “lytic”. A water that meets all three of these collective variable conditions is positioned in a small volume of the NTPase active site, is hydrogen bonding with the active site’s proton acceptor (Glu285) (data not shown), and the water’s dipole moment vector is approximately facing the terminal phosphoanhydride bond vector, as proposed in the SN2 mechanism for the hydrolysis reaction.

The projected covariance magnitude represents the fluctuation between G414 (source residue) and S364 and K366 (sink residues) projected onto the vector spanning the helical gate between subdomain 2 and subdomain 3 of NS3h. G414 was chosen as the source residue due to its coordination with the lytic water within the ATP binding pocket(24), and S364 and K366 were chosen as the sink residues due to their location in the helical gate and large magnitude covariance with G414. The covariance tensors between G414-S364 and G414-K366 were then dotted into the helical gate open/closing vector to obtain how the fluctuation perturbs the opening/closing of the helical gate.

## ACKNOWLEDGEMENTS

We would like to acknowledge the support of NVIDIA Corporation with the donation of one of the Titan X Maxwell GPUs used for this research. This work was supported by NIH grant R01 Al132668 to BJG and XSEDE resources under the allocation CHE160008 as well as CSU funding to MM. We would also like to acknowledge the help of Dr. Christopher Berndsen in analyzing the kinetic data, as well as the helpful discussions with Dr. Olve Peersen for analyzing the kinetic data, and any other helpful discussions with members of the Geiss and McCullagh labs.

## AUTHOR CONTRIBUTIONS

BJG, MM, and KED conceived of the project. KED and RBD performed molecular dynamics simulations and analysis. KED performed all experimental aspects of the project. MM assisted with computational analyses, and BJG assisted with experimental analyses. MM and BJG oversaw the research. KED wrote the manuscript, and MM and BJG revised the manuscript with input from all authors.

