## Supplemental Data for "Motif V acts as a Regulator of Energy Transduction Between the Flavivirus NS3 ATPase and RNA Binding Cleft"

**Kelly E. Du Pont[1,2], Russell Davidson[1], Martin McCullagh[1,4*], Brian J. Geiss[2,3,4*]**
From the [1]Department of Chemistry, Colorado State University, Fort Collins CO 80523; [2]Department of Microbiology, Immunology, and Pathology, Colorado State University, Fort Collins, CO 80523; [3]School of Biomedical Engineering, Colorado State University, Fort Collins, CO 80523

Figure S1

Figure S2

Table S1

Table S2

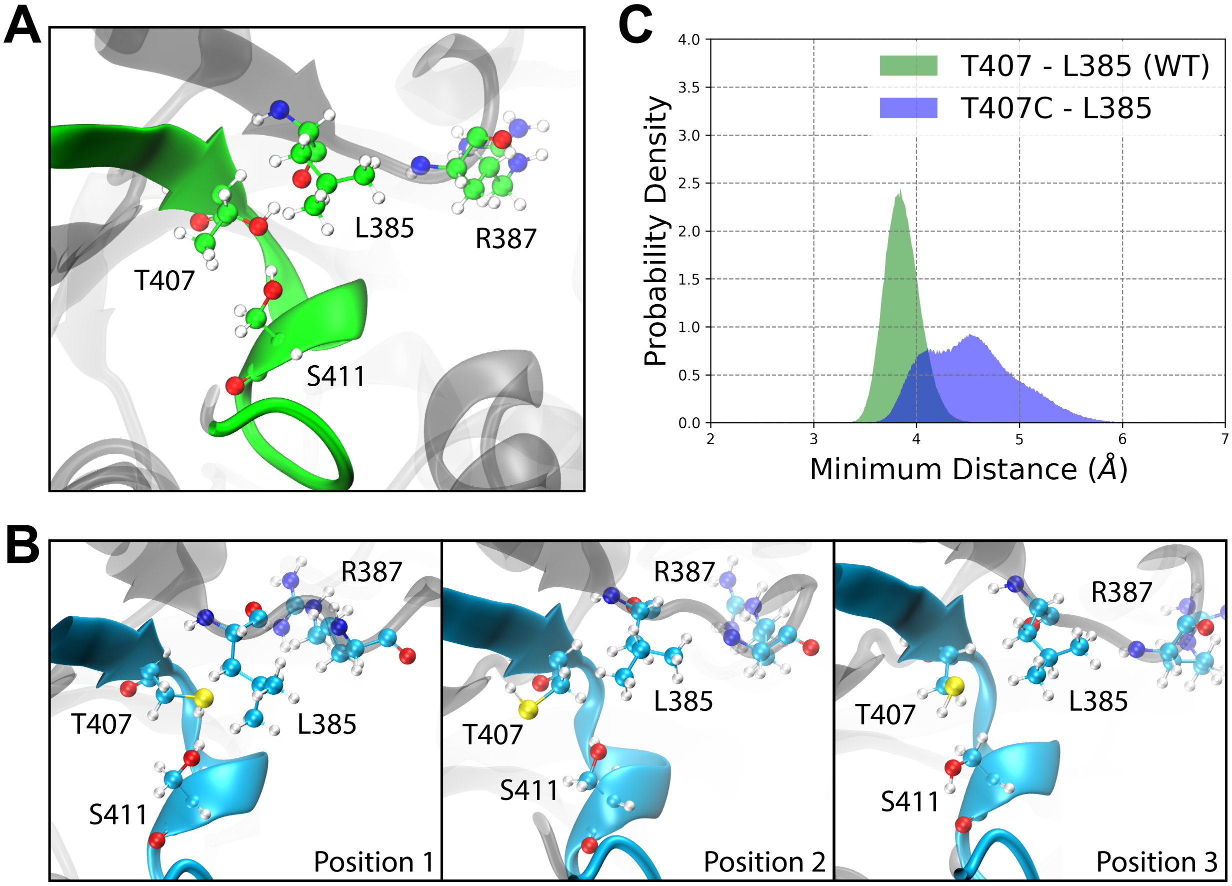


**Figure S1. Methyl group in the T407 sidechain stabilizes T407 interactions with L385. A)** The methyl group stabilizes the structure of T407 in the WT simulations. **B)** Without the methyl group, the sulfur group of T407C fluctuates frequently throughout the T407C simulations. **C)** The minimum distance was calculated between the L387 sidechain and either the hydroxyl group (threonine sidechain) or the thiol group (cysteine sidechain) at position 407 for each replicate of the WT and the T407C simulations. The probability density of the average minimum distance was then determined for WT and T407C simulations.

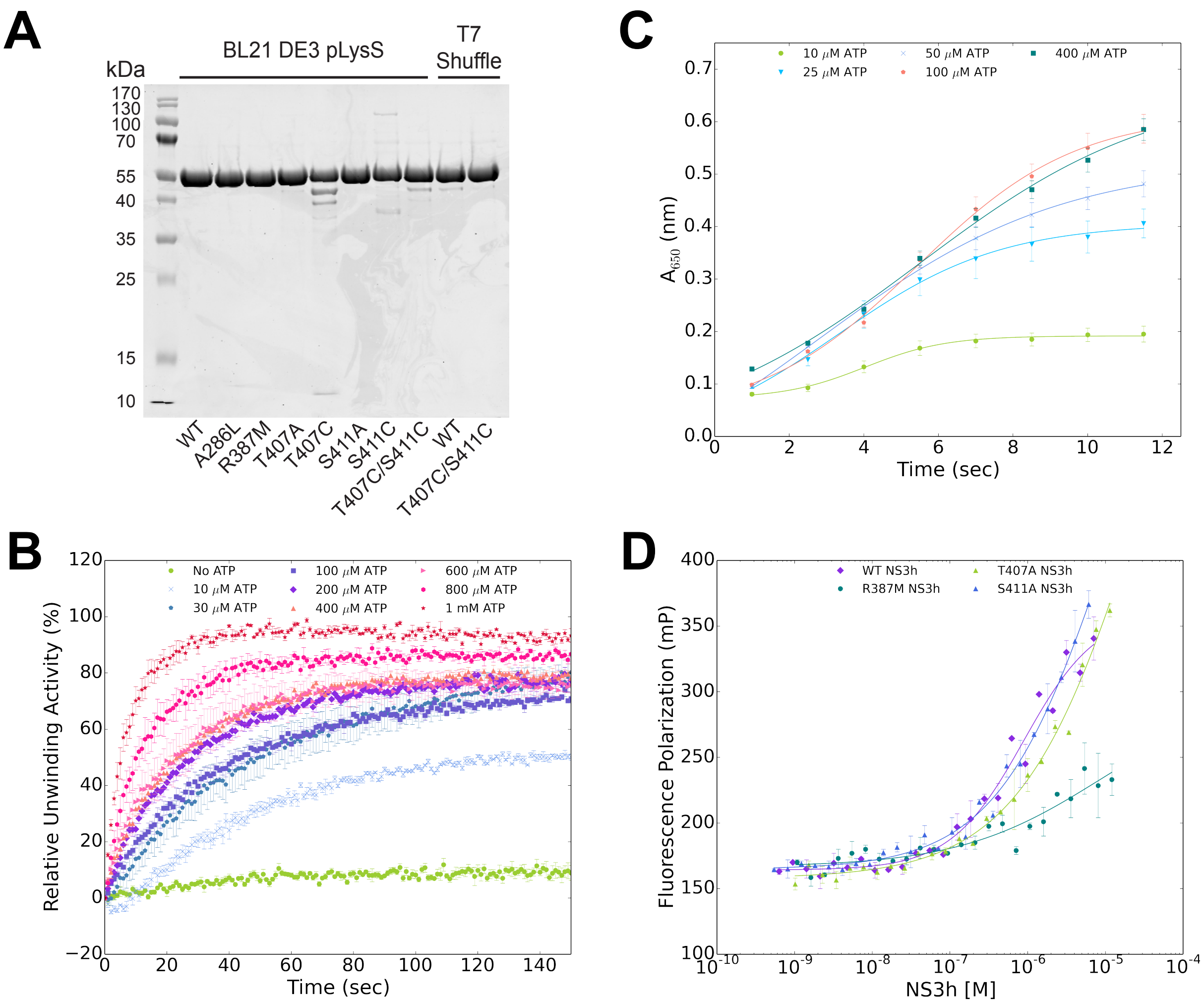


**Figure S2. Confirmation and experimental design of biochemical assays. A)** Recombinant NS3 helicase variants were purified and confirmed on SDS-PAGE. We noted that T407C degraded over time but it was full-length when used in assays. **B)** RNA binding affinity of NS3h were measured using fluorescence polarization of a single-stranded RNA (52). **C)** ATPase activity was measured using BIOMOL Green absorbance at 650 nm over time. **D)** RNA helicase activity was measured by increase of fluorescence over time.

| Virus | Number of Sequences | 404 Consensus | F404(%) | Y404(%) | L404(%) | 405 Consensus | V405(%) | L405(%) | I405(%) | 406 Consensus | V406(%) | D406(%) | G406(%) | I406(%) | L406(%) | 407 Consensus | T407(%) | A407(%) | S407(%) | 411 Consensus | S411(%) | A411(%) |
|---|---|---|---|---|---|---|---|---|---|---|---|---|---|---|---|---|---|---|---|---|---|---|
| Karshi Virus | 1 | F | 100.0 | - | - | V | 100.0 | - | - | V | 100.0 | - | - | - | - | T | 100.0 | - | - | S | 100.0 | - |
| Powassan | 22 | F | 100.0 | - | - | V | 100.0 | - | - | V | 100.0 | - | - | - | - | T | 100.0 | - | - | S | 100.0 | - |
| MMLV | 3 | F | 100.0 | - | - | I | - | - | 100.0 | L | - | - | - | - | 100.0 | T | 100.0 | - | - | S | 100.0 | - |
| Rio Bravo | 3 | F | 100.0 | - | - | I | - | - | 100.0 | L | - | - | - | - | 100.0 | T | 100.0 | - | - | S | 100.0 | - |
| Modoc | 3 | F | 100.0 | - | - | I | - | - | 100.0 | L | - | - | - | - | 100.0 | T | 100.0 | - | - | S | 100.0 | - |
| Apoi | 2 | F | 100.0 | - | - | I | - | - | 100.0 | L | - | - | - | - | 100.0 | T | 100.0 | - | - | S | 100.0 | - |
| JEV | 321 | F | 100.0 | - | - | V | 100.0 | - | - | I | 0.9 | - | - | 99.1 | - | T | 100.0 | - | - | S | 100.0 | - |
| Usutu | 138 | F | 100.0 | - | - | V | 100.0 | - | - | I | 0.7 | - | - | 99.3 | - | T | 100.0 | - | - | S | 100.0 | - |
| WNV | 2008 | F | 100.0 | - | - | V | 100.0 | - | - | I | 0.6 | - | - | 99.4 | - | T | 100.0 | - | - | S | 100.0 | - |
| Kunjin | 44 | F | 100.0 | - | - | V | 100.0 | - | - | V | 100.0 | - | - | - | - | T | 100.0 | - | - | S | 100.0 | - |
| Bagaza | 12 | F | 100.0 | - | - | V | 100.0 | - | - | I | - | - | - | 100.0 | - | T | 100.0 | - | - | S | 100.0 | - |
| Ilheus | 1 | F | 100.0 | - | - | V | 100.0 | - | - | I | - | - | - | 100.0 | - | T | 100.0 | - | - | S | 100.0 | - |
| Kedougou | 1 | F | 100.0 | - | - | V | 100.0 | - | - | I | - | - | - | 100.0 | - | T | 100.0 | - | - | S | 100.0 | - |
| Zika | 659 | F | 100.0 | - | - | V | 100.0 | - | - | I | 85.6 | - | - | 14.4 | - | T | 100.0 | - | - | S | 100.0 | - |
| Bussuquara | 1 | F | 100.0 | - | - | V | 100.0 | - | - | V | 100.0 | - | - | - | - | T | 100.0 | - | - | S | 100.0 | - |
| DENV1 | 1948 | Y | 0.1 | 99.9 | - | V | 100.0 | - | - | V | 99.9 | - | 0.1 | - | - | T | 100.0 | - | - | S | 100.0 | - |
| DENV2 | 1524 | F | 100.0 | - | - | V | 100.0 | - | - | V | 99.8 | 0.1 | - | 0.1 | - | T | 100.0 | - | - | S | 100.0 | - |
| DENV3 | 1112 | F | 100.0 | - | - | V | 99.9 | 0.1 | - | V | 99.9 | - | 0.1 | - | - | T | 99.8 | 0.2 | - | S | 100.0 | - |
| DENV4 | 241 | F | 100.0 | - | - | V | 100.0 | - | - | V | 100.0 | - | - | - | - | T | 100.0 | - | - | S | 100.0 | - |
| Kokobera | 1 | F | 100.0 | - | - | V | 100.0 | - | - | I | - | - | - | 100.0 | - | T | 100.0 | - | - | S | 100.0 | - |
| YFV | 105 | F | 100.0 | - | - | I | - | - | 100.0 | L | - | - | - | - | 100.0 | A | - | 100.0 | - | A | - | 100.0 |
| Sepik | 1 | F | 100.0 | - | - | I | - | - | 100.0 | L | - | - | - | - | 100.0 | A | - | 100.0 | - | A | - | 100.0 |
| Entebbe Bat | 1 | F | 100.0 | - | - | I | - | - | 100.0 | L | - | - | - | - | 100.0 | T | 100.0 | - | - | A | - | 100.0 |
| Cell fusing agent | 35 | L | - | - | 100.0 | V | 100.0 | - | - | I | - | - | - | 100.0 | - | S | - | - | 100.0 | S | 100.0 | - |
| Kamiti River | 2 | L | - | - | 100.0 | V | 100.0 | - | - | V | 100.0 | - | - | - | - | S | - | - | 100.0 | S | 100.0 | - |
| Culex | 32 | F | 100.0 | - | - | I | - | - | 100.0 | V | 100.0 | - | - | - | - | S | - | - | 100.0 | S | 100.0 | - |

**Table S1. Flavivirus sequence variability at positions 404, 405, 406, 407 and 411.** Flavivirus sequences were analyzed to determine the consensus sequence at motif V positions 404, 405, 406, 407 and 411. The percentage of finding the consensus or another residue instead of the consensus are reported for each position for each flavivirus.

### Substate Inhibition of ATPase Activity

| NS3h variant | $K_i$ (μM) |
|---|---|
| WT | 888.2 ± 247.7 |
| A286L | 570.5 ± 377.4 |
| R387M | 745.7 ± 371.1 |
| T407A | 1442.1 ± 727.5 |
| T407C | 587.6 ± 177.8 |
| S411A | 695.5 ± 404.4 |
| S411C | 775.8 ± 166.9 |
| T407C/S411C | 1157.8 ± 480.1 |
| WT reduced† | 534.3 ± 110.5 |
| WT oxidized‡ | 297.3 ± 133.0 |
| T407C/S411C reduced† | 729.0 ± 148.4 |
| T407C/S411C oxidized‡ | 532.7 ± 137.3 |

**Table S2. ATP substrate inhibition of ATPase activity.** The ATPase activity exhibited substrate inhibition at high concentrations of ATP. As a result of substrate inhibition, the data was fit to the substrate inhibition equation. The inhibition constant ($K_i$) was reported for each NS3 helicase variant.
